# PTENP1-AS contributes to BRAF inhibitor resistance and is associated with adverse clinical outcome in stage III melanoma

**DOI:** 10.1101/2020.05.03.073627

**Authors:** Linda Vidarsdottir, Alireza Azimi, Ingibjorg Sigvaldadottir, Aldwin Suryo Rahmanto, Andreas Petri, Sakari Kauppinen, Christian Ingvar, Göran Jönsson, Håkan Olsson, Marianne Frostvik Stolt, Rainer Tuominen, Olle Sangfelt, Katja Pokrovskaja Tamm, Johan Hansson, Dan Grandér, Suzanne Egyházi Brage, Per Johnsson

## Abstract

Approximately 50% of human cutaneous melanomas carry activating mutations in the serine/threonine protein kinase BRAF. BRAF inhibitors (BRAFi) selectively target the oncogenic BRAF^V600E/K^ and are effective in approximately 80% of patients carrying the mutation. However, resistance to BRAFi is common and emerges within a median time of 6-7 months of treatment and is prolonged to 11 months when combined with MEK inhibitors. Better characterization of the underlying molecular processes is therefore needed to further improve treatments. Inactivation of the tumor suppressor gene *PTEN* has been suggested to occur during melanomagenesis and drug resistance development. We recently demonstrated that transcription of *PTEN* is negatively regulated by an antisense RNA from the PTEN pseudogene (PTENP1-AS) and here set out to investigate the impact of this molecular pathway on the resistance to BRAFi and clinical outcome. We used a panel of BRAFi resistant A375 sublines, and observed increased levels of PTENP1-AS associated with reduced expression of *PTEN*. Furthermore, this loss of *PTEN* expression was correlated to increased recruitment of Enhancer of zeste homolog 2 (EZH2) and formation of the transcriptional repression mark H3K27me3 at the *PTEN* promoter in the resistant cells. We demonstrated that targeting of PTENP1-AS was able to re-activate the expression of *PTEN* and sensitize resistant melanoma cells to BRAFi. Finally, we showed that PTENP1-AS is a promising prognostic marker for clinical outcome in melanoma patients as high expression of PTENP1-AS in regional lymph node metastases from stage III melanoma patients correlated with poor survival.

## BACKGROUND

Although cutaneous malignant melanoma is a molecularly diverse disease, approximately 50% carry activating mutations in the serine/threonine protein kinase BRAF. The majority of BRAF mutations are represented by a valine (V) to glutamic acid (E) or lysine (K) substitution at position 600 (BRAF^V600E/K^)^1^. These missense mutations occur in the BRAF kinase domain, which results in constitutively active BRAF and subsequently activated MAPK signaling. Recent drug development efforts using a targeted approach have successfully led to the development of small inhibitory molecules (BRAFi), which efficiently and specifically target the oncogenic BRAF^V600E/K^. Vemurafenib represents one such inhibitor and 80% of patients with advanced melanoma carrying a BRAF^V600E/K^ mutation initially respond well to this treatment and show tumor regression^2^. However, the development of resistance to BRAFi is a major issue and emerges within a median time of 6-7 months of treatment which can be prolonged to 11 months when combined with MEK inhibitors. The 5-year overall survival rate is only 34% for the combination treatment^3^. In recent years there has been a growing understanding of the underlying molecular mechanisms involved in acquired BRAFi resistance^4,5^. Activation of the PI3K/AKT pathway due to loss of PTEN has been shown to be one of the mechanisms contributing to BRAFi resistance^6^. Thus, dissecting the mechanisms of PTEN loss is vital in order to design novel approaches for treatment, which may circumvent or delay the onset of resistance.

Approximately 14,000 pseudogenes have been identified in the human genome and hundreds of those shown to be transcribed into non-coding RNAs (ncRNAs)^7^. Transcribed pseudogenes have been suggested to be capable of acting as regulators of gene expression^8–10^, for instance through mechanisms such as microRNA (miRNA) sponging and *trans* acting antisense RNA (asRNA) regulation. One of these, the PTEN pseudogene, PTENP1 (also known as PTENpg1, PTENΨ), has been shown to be transcribed into both sense and asRNA transcripts. While the sense transcript (PTENP1-S) functions as a positive regulator of *PTEN* by acting as a miRNA sponge for *PTEN* related miRNAs^10^, the asRNA transcript (PTENP1-AS) functions as a negative regulator of *PTEN* expression through inducing epigenetic alterations at the *PTEN* promoter^8^. *PTEN* is a tumor suppressor gene which is frequently inactivated across a wide range of cancers such as breast cancer, prostate cancer and melanoma (reviewed in^11^). Loss of PTEN expression has been reported to play a role not only in the development of resistance to BRAFi^6^, but also in the process of melanoma metastasis^12^. Moreover, mutated *BRAF* itself is not sufficient to induce melanoma, other molecular events such as inactivation of *PTEN*^12,13^, are required for melanoma initiation.

In this study, our aim was to study the involvement of PTENP1-AS in melanoma tumors and in resistance to the BRAFi vemurafenib. We observe that the expression of PTENP1-AS is induced in BRAFi resistant cell lines, which coincides with transcriptional suppression of *PTEN*. We further show that targeting of the PTENP1-AS transcript sensitizes resistant melanoma cells to vemurafenib. Mechanistically, we conclude that up-regulation of the transcription factor C/EBPβ initiates transcription of PTENP1-AS and that *PTEN* is suppressed through the recruitment of EZH2 to the *PTEN* promoter with subsequent formation of H3K27me3. Finally, we also demonstrate that expression of PTENP1-AS is increased in tumor samples from stage III melanoma patients with poor survival.

## RESULTS

### Inverse correlation of *PTEN* and PTENP1-AS in BRAFi resistant A375 sublines

Previous studies suggested downregulation of *PTEN* to be an important mechanism for the development of resistance to BRAFi^6,12^. Based on these reports, we hypothesized that the PTENP1-AS transcript could be involved in this process through transcriptional regulation of *PTEN*. To study the involvement of PTENP1-AS, a series of BRAFi resistant melanoma cell lines were obtained by culturing the BRAFi sensitive A375 cell line had been cultured in increasing doses of BRAFi (A375PR1 (resistant to PLX4720), A375VR3 and A375VR4 (both resistant to vemurafenib)^14^. Upon IC50 measurements, all three cell lines displayed at least 10 times increased tolerance to the BRAFi vemurafenib compared to the parental A375 cells. Our results demonstrated downregulation of PTEN at the protein levels in all three resistant sublines compared to the A375 parental cell line while the negatively regulated downstream target of PTEN, p-AKT, showed increased phosphorylation in all resistant A375 sublines (**Figures 1A, S1A**). Downregulation of *PTEN* was also confirmed at the mRNA level in all three resistant sublines compared to the parental A375 cell line (**Figures 1B, S1B**). In summary, these initial observations motivated us to further elucidate the underlying molecular events mediating suppression of PTEN.

**Figure 1.**
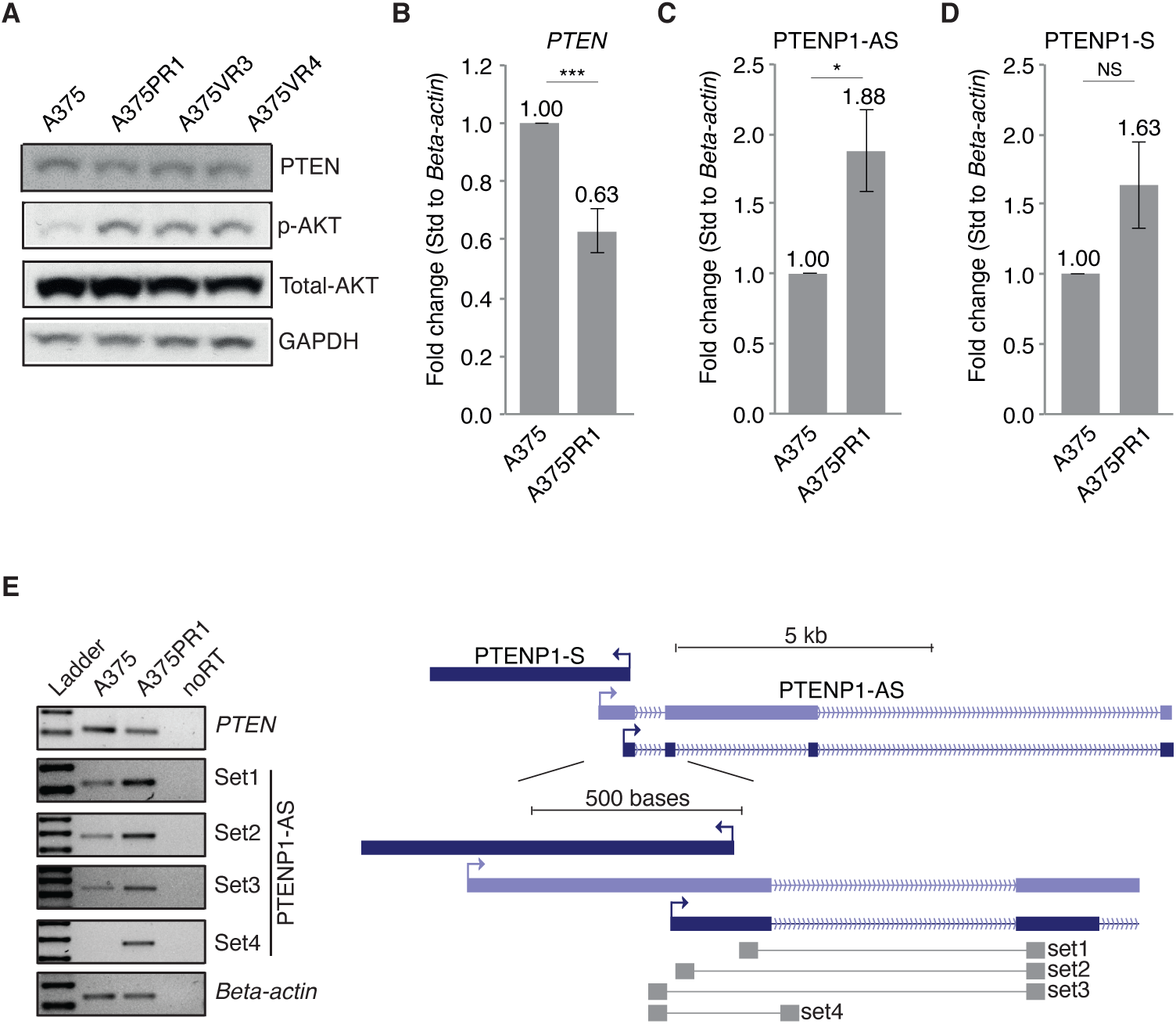
Expression of PTEN and PTENP1-AS in A375 and vemurafenib resistant sublines. **(A)** Western blot analysis presenting the expression of PTEN, total AKT and p-AKT in the A375 and BRAFi resistant sublines. **(B-D)** qRTPCR analysis of **(B)** *PTEN,* **(C)** PTENP1-AS and **(D)** PTENP1-S in A375 and the BRAFi resistant A375PR1 subline (n=3, p-values represent a two-tailed student’s t-test). **(E)** Semi-qRTPCR assay showing different isoforms of the PTENP1-AS transcripts with a scheme depicting primer binding sites.

We next sought to determine the involvement of PTENP1 encoded lncRNAs in this process. To this end, we measured the levels of sense and antisense transcripts and found PTENP1-AS levels to be significantly upregulated in all three resistant sublines (**Figures 1C, S1C**). Notably, we also observed a modest, although non-significant, induction of PTENP1-S (**Figures 1D, S1D**). This is, however, in contrast to the previously reported miRNA-sponge function of PTENP1-S, since such a mechanism would result in increased expression of *PTEN* due to the release of miRNA-mediated suppression of *PTEN*. We therefore considered PTENP1-AS to hold the main regulatory function under these conditions.

Previously, we have reported multiple isoforms of PTENP1-AS with distinct regulatory functions^8^. We selected the A375PR1 subline for further characterization of isoforms using a total of four different primer sets. A concurrent induction was observed for all primer sets (**Figure 1E**). Notably, the level of unspliced form of PTENP1-AS was found to be induced in the resistant cells, indicating that induction of PTENP1-AS takes place at the transcriptional level.

### PTENP1-AS suppresses *PTEN* in vemurafenib resistant cells through chromatin remodelling

To investigate the molecular interplay between PTENP1-AS and *PTEN* in greater detail, we next designed a gapmer antisense oligonucleotide (ASO) (**Figures 2A-B**) as well as an siRNA targeting the PTENP1-AS transcript (**Figures 2C-D**). Both approaches decreased the amounts of PTENP1-AS with more than 60% in A375 as well as in A375PR1 cells. Intriguingly, knockdown of PTENP1-AS only induced the expression of *PTEN* in the A375PR1 cells, while *PTEN* remained unaffected in the vemurafenib sensitive A375 cells (**Figures 2A-D**), suggesting that PTENP1-AS is only involved in suppression of *PTEN* in the resistant cells.

**Figure 2.**
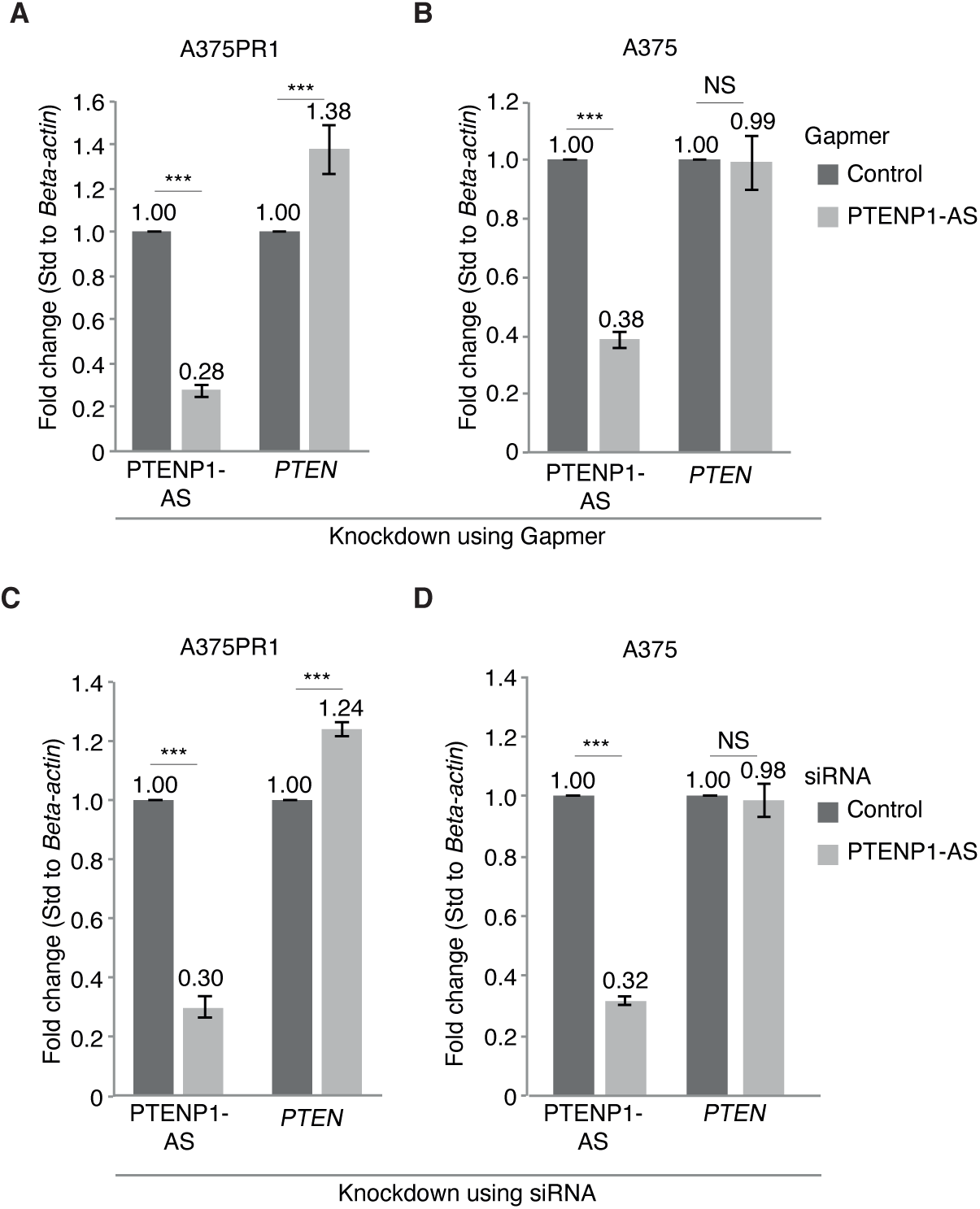
Knockdown of PTENP1-AS activates PTEN in vemurafenib resistant cells. **(A)** qRTPCR measuring the expression of PTENP1-AS and *PTEN* in A375PR1 cells upon knockdown of PTENP1-AS using gapmer ASO. **(B)** qRTPCR measuring the expression of PTENP1-AS and *PTEN* in A375 cells upon knockdown of PTENP1-AS using gapmer ASO. **(C)** qRTPCR measuring the expression of PTENP1-AS and *PTEN* in A375PR1 cells upon knockdown of PTENP1-AS using siRNA. **(D** qRTPCR measuring the expression of PTENP1-AS and *PTEN* in A375 cells upon knockdown of PTENP1-AS using siRNA (A-D; n=3, p-values represent a two-tailed student’s t-test)

To further investigate the function of PTENP1-AS, we next asked if the resistant A375PR1 subline had acquired epigenetic changes at the *PTEN* promoter. We first assessed DNA methylation using the McrBc enzyme that cleaves methylated DNA at Pu^m^CG sequence elements. Elevated DNA methylation was demonstrated by increased enzymatic cleavage at the *PTEN* promoter in the resistant A375PR1 subline compared to parental A375 cells (**Figure 3A**). Since PTENP1-AS has previously been shown to recruit the Enhancer of zeste homolog 2 (EZH2) and DNA methyltransferase 3a (DNMT3A) to the *PTEN* promoter^8^, we next investigated the involvement of these factors upon development of resistance. Although individual siRNA-induced knockdown of *DNMT3a* and *EZH2* (**Figures S2A-B**) only resulted in a modest induction of *PTEN* (**Figures 3B-C**), the combined knockdown generated over a twofold induction of *PTEN* in the A375PR1 cells (**Figures S2C-D)**, while the expression of *PTEN* was largely unaffected in the parental A375 cells (**Figure 3D**). We also confirmed the baseline levels of *DNMT3A, EZH2* and H3K27me3, the EZH2 downstream target, to be similar in the resistant and parental cells (**Figures 4A-C**), suggesting that suppression of *PTEN* is not an indirect consequence of genome-wide chromatin changes. These observations prompted us to look more specifically for EZH2 and H3K27me3 levels at the *PTEN* promoter. The resistant A375PR1 cells showed enrichment of EZH2 (**Figure 4D**) and increased presence of the transcriptional suppressive chromatin mark H3K27me3 at the *PTEN* promoter (**Figure 4E**). Taken together, these data suggest that suppression of *PTEN* is mediated by PTENP1-AS through the recruitment of chromatin remodeling factors utilizing a mechanism that may be exclusively active in the vemurafenib resistant cells.

**Figure 3.**
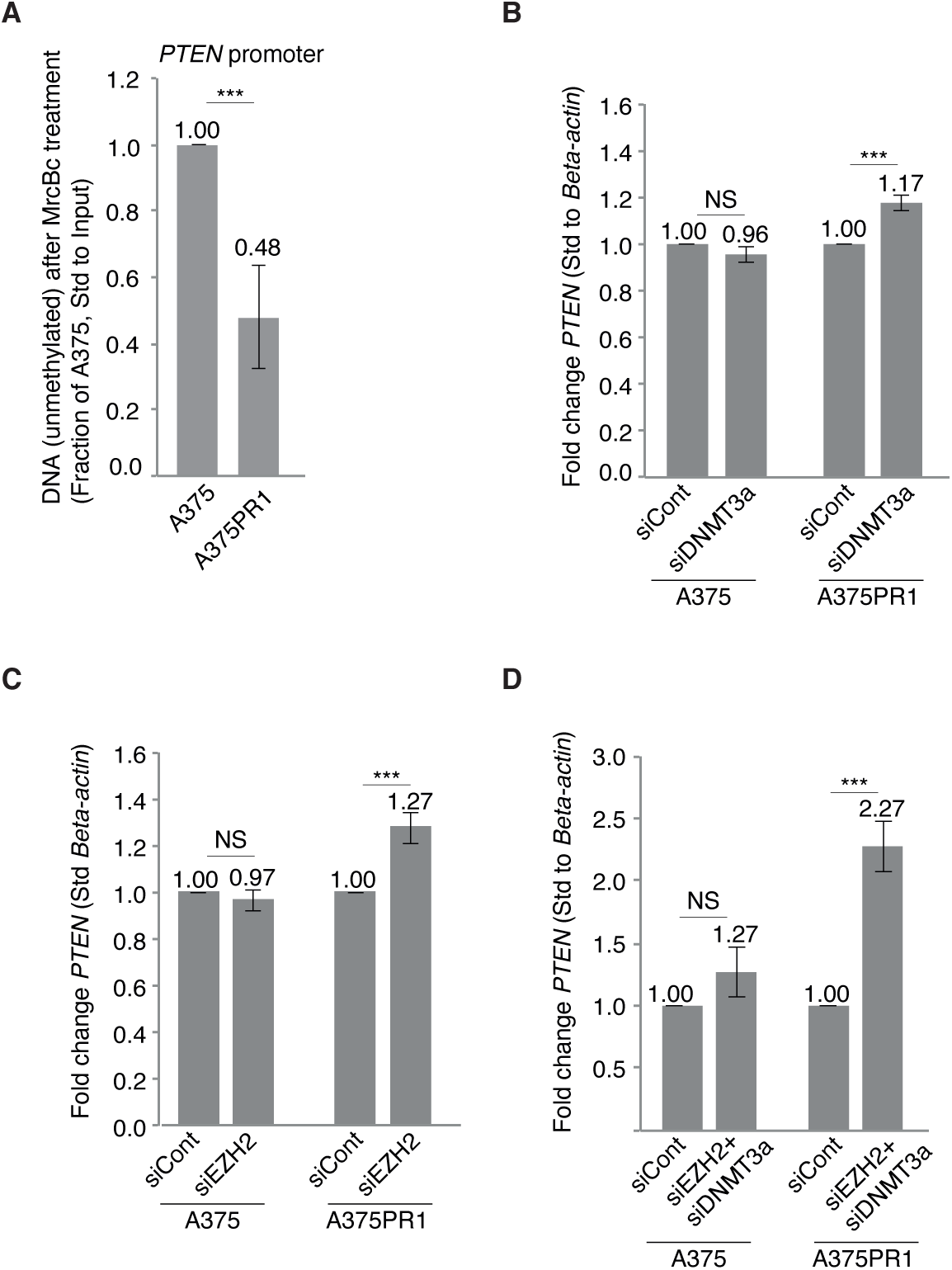
Knockdown of EZH2 and DNMT3A induces the expression of PTEN. **(A)** qRTPCR measuring the remaining (unmethylated) DNA of the *PTEN* promoter after MrcBc cleavage (n=3). **(B)** qRTPCR of *PTEN* following siRNA induced knockdown of *DNMT3A* in A375 and A375PR1 cells (n>3). **(C)** Expression of *PTEN* by qRTPCR following knockdown of *EZH2* in A375 and A375PR1 cells (n>3). **(D)** Expression of *PTEN* by qRTPCR following co-knockdown of *EZH2* and *DNMT3a* in A375 and A375PR1 cells (n=3) (A-D; p-values represent a two-tailed student’s t-test).

**Figure 4.**
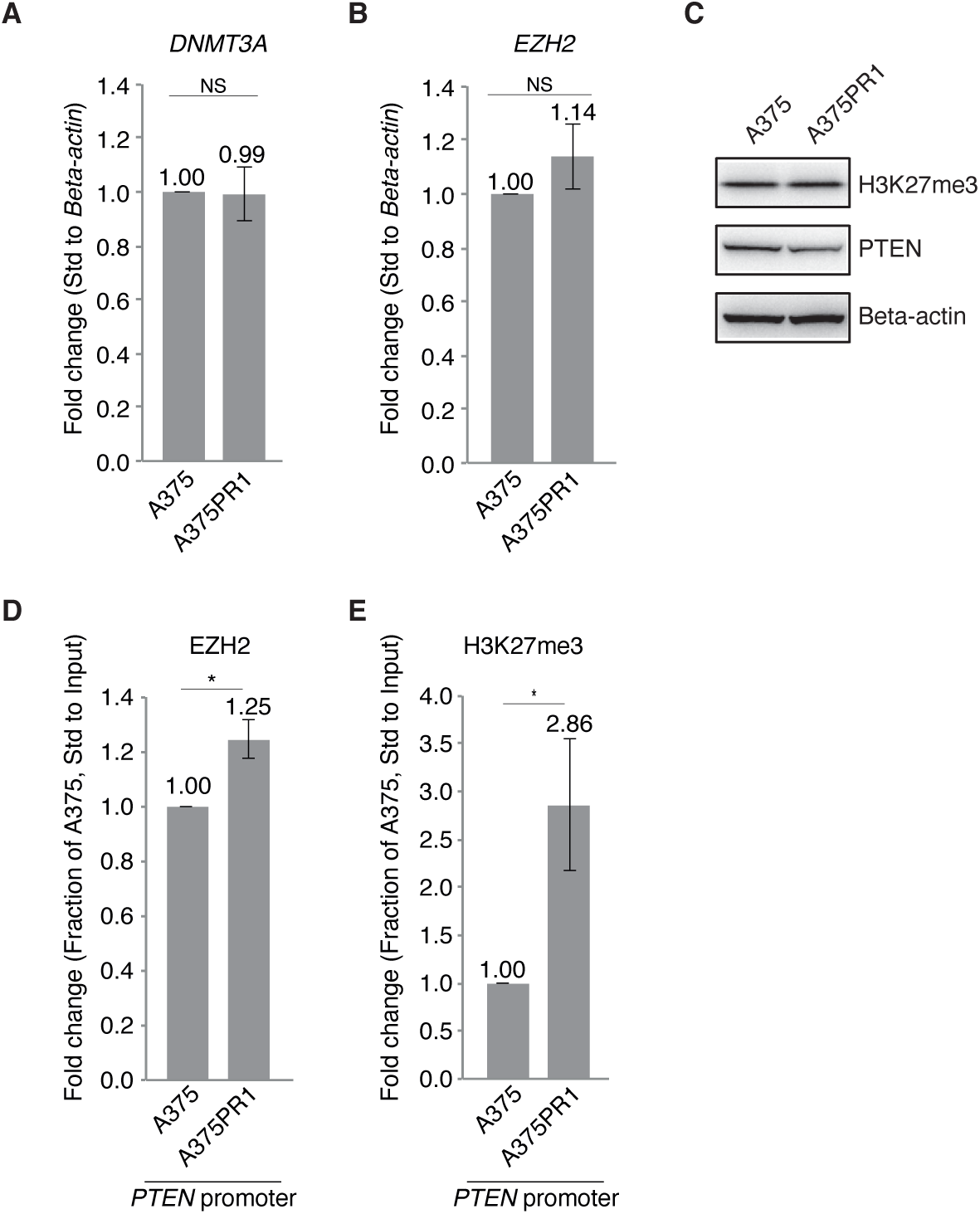
Suppression of PTEN is mediated by EZH2 and DNMT3A. **(A)** qRTPCR showing expression levels of *DNMT3A* in the A375 and A375PR1 cells (n=3). **(B)** qRTPCR showing expression levels of *EZH2* in the A375 and A375 PR1 cells (n=3). **(C)** Western blot measuring the levels of H3K27me3 in A375 and A375PR1 cells. **(D)** ChIP analysis assessing the levels of EZH2 at the *PTEN* promoter in the A375 and A375PR1 cells (n=3). **(E)** ChIP analysis assessing the levels of H3K27me3 at the *PTEN* promoter in the A375 and A375PR1 cells (n=3).

### C/EBPβ is a transcriptional regulator of PTENP1-AS

To explore the molecular events underlying transcriptional induction of PTENP1-AS in the BRAFi resistant sublines, we took advantage of FANTOM5 CAGE data and ChIP-seq data from the UCSC Genome Browser, and identified enrichment of the transcription factor C/EBPβ upstream of the PTENP1-AS transcriptional start site (TSS) (**Figure 5A**). C/EBPs can homodimerize or heterodimerize and bind to the consensus DNA sequence RTTGCGYAAY^15^, (R is an A/G, Y is C/T). We identified a binding motif for C/EBPs upstream of the PTENP1-AS TSS (**Figure 5A**) and also found the expression of *C/EBPβ* mRNA to be induced in the BRAFi resistant cells (**Figure 5B**). A ChIP pulldown confirmed enrichment of C/EBPβ at the PTENP1-AS promoter as compared to the BRAFi sensitive cells (**Figure 5C**). Finally, siRNA-mediated knockdown of *C/EBPβ* reduced the expression of unspliced PTENP1-AS in the A375PR1 cells, while the expression of PTENP1-S was not affected (**Figure S3A**). In summary, these observations indicate that C/EBPβ functions as a transcriptional regulator of PTENP1-AS.

**Figure 5.**
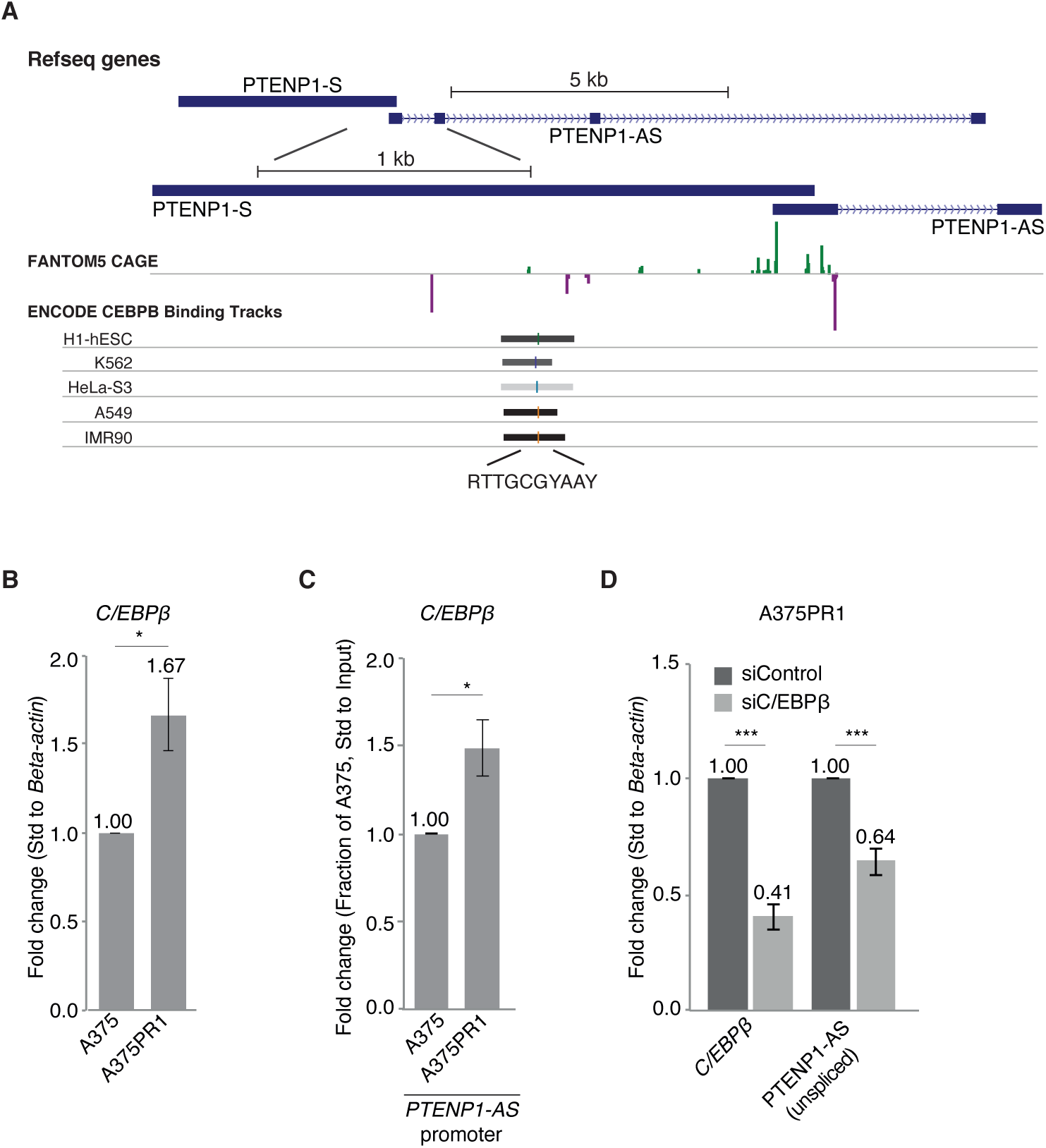
C/EBPβ is a transcriptional regulator of PTENP1-AS. **(A)** A scheme showing enrichment of CEBPβ at the PTENP1 locus (data retrieved from the Zenbu FANTOM5 CAGE and the UCSC genome browser) and the predicted DNA binding site for *C/EBPβ*. **(B)** qRTPCR assessing the expression of *C/EBPβ* in A375 and A375PR1 cells (n>3). **(C)** ChIP assesing *C/EBP*β binding to the PTENP1-AS promoter in the A375 and A375PR1 cells (n=3). **(D)** qRTPCR assessing the expression of *C/EBP*β and unspliced PTENP1-AS in A375PR1 cells upon knockdown of *C/EBPβ* (n=3).

### PTENP1-AS is a clinically relevant target and a prognostic marker in melanoma

We next explored if the resistant A375PR1 cells could be sensitized to BRAFi by targeting the PTENP1-AS transcript. Notably, gapmer ASO-induced knockdown of PTENP1-AS resulted in increased induction of apoptosis to 10 μM of vemurafenib in the A375PR1 cells (**Figure 6A**).

**Figure 6.**
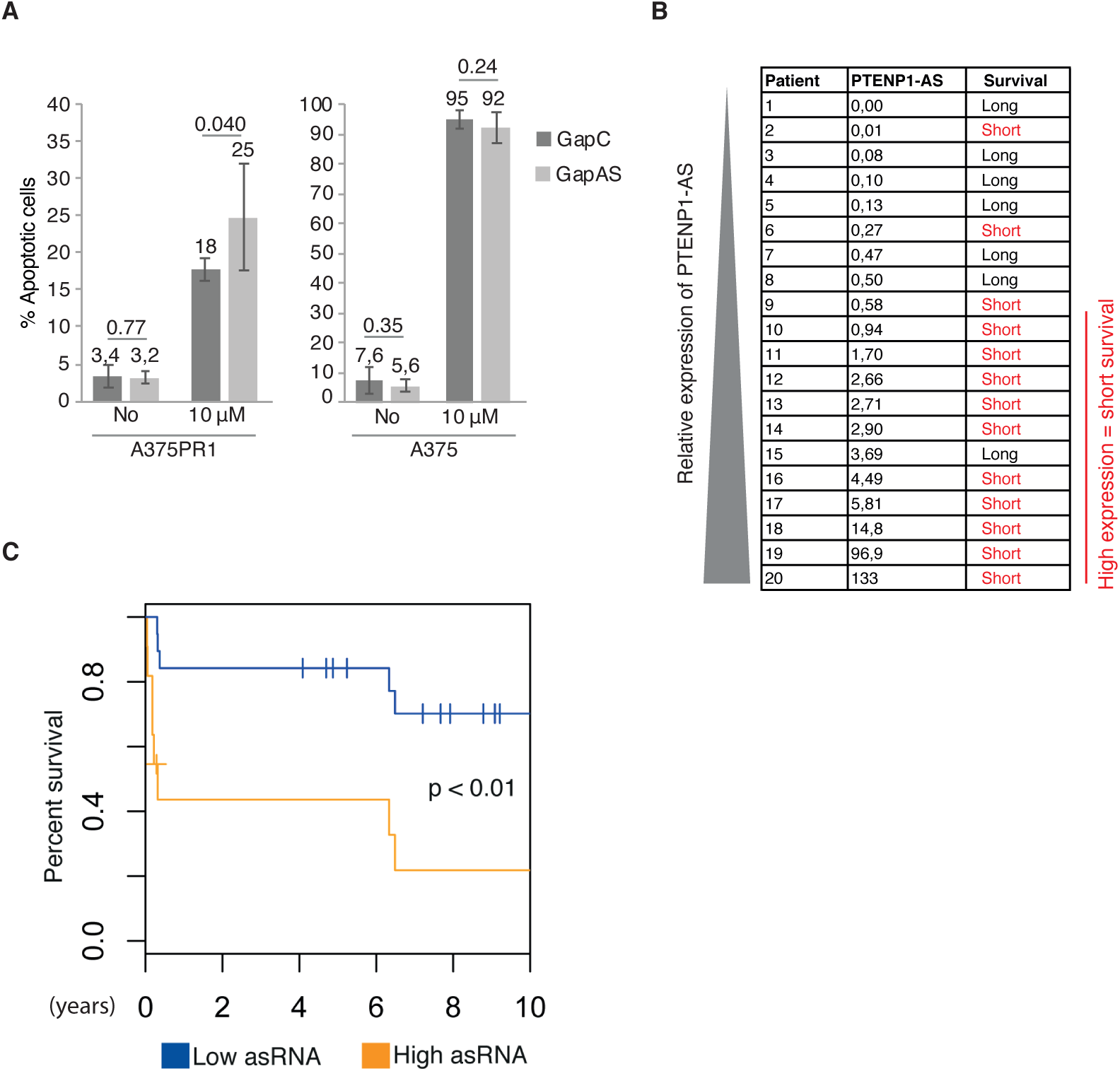
Manipulation of the PTENP1-AS pathway resensitizes resistant A375PR1 sub line to vemurafenib treatment. **(A)** Apoptotic cells after knockdown of PTENP1-AS upon vemurafenib treatment (n=6). **(B)** qRTPCR presenting the expression of PTENP1-AS in a set of 20 first regional lymph node metastases from stage III melanoma patients. Based on clinical follow up data, the patients were categorized as long- or short-term survivors, >60 months or ≤13 months. **(C)** The expression of PTENP1-AS was determined in an independent set of 29 stage III melanoma patients with first regional lymph node metastases. The patients were divided according to high (n=12) or low (n=17) expression of PTENP1-AS and overall survival was analyzed in a Kaplan-Meier plot. The p-value represents a log-rank test.

We finally set out to investigate the relevance of PTENP1-AS for clinical outcome in melanoma. The expression of PTENP1-AS was first determined in lymph node metastases from a cohort of 20 stage III melanoma patients prior to the treatment. The samples were initially chosen to include two groups of patients, with either long (≥ 60 months) or short (≤13 months) overall survival^16^, and the expression of PTENP1-AS was determined by qRTPCR. Notably, 9/10 (90%) patients having tumors with high expression of PTENP1-AS (median cut-off) had poor survival, while low expression of PTENP1-AS seemed to be indicative of prolonged survival, although not reaching significance (p=0.06) (**Figure 6B**). This motivated us to evaluate the expression of PTENP1-AS in an independent second cohort of stage III melanoma samples. Lymph node metastases from 29 stage III melanoma patients were analyzed for the expression of PTENP1-AS. The patient samples from the second cohort were divided in high (n=12) or low (n=17) expression of PTENP1-AS, as determined by qRTPCR, and a survival plot was generated (**Figure 6C**). A significant difference in overall survival was observed between patients with high and low expression of PTENP1-AS. In summary, the expression of PTENP1-AS appears to be a promising prognostic marker for clinical outcome, where increased expression of PTENP1-AS correlates with poor overall survival.

## DISCUSSION

The data presented in this study suggest a role for the PTENP1-AS in the development of resistance to BRAFi. On the basis of our data, we present a hypothesis, where the PTENP1-AS transcript is induced by the transcription factor C/EBPβ in BRAFi resistant melanoma cell lines, which results in transcriptional suppression of *PTEN* through the recruitment of EZH2 to the *PTEN* promoter and subsequent formation of H3K27me3 and DNA methylation. Moreover, we find that expression of PTENP1-AS predicts clinical outcome in stage III melanoma patients, where high expression of PTENP1-AS in first regional lymph node metastases correlates with poor overall survival. In summary, our findings bring important information about underlying molecular events involved in the development of BRAFi resistance and also present PTENP1-AS as a promising prognostic marker for clinical outcome of stage III melanoma. Particularly, by gaining insight for this pathway in more detail, we could develop novel approaches to resensitize drug resistant cells.

Our data are consistent with previous findings demonstrating that the PTENP1-AS is a negative regulator of *PTEN* through chromatin remodeling of the *PTEN* promoter^8^. Targeting the PTENP1-AS transcript using gapmer ASOs and siRNAs generates a moderate induction of *PTEN* exclusively in the resistant subline, thus demonstrating that the PTENP1-AS transcript does not act as an on/off switch for *PTEN* expression, but rather causes consistent but subtle variations of transcription in a phenotype-dependent manner^8^. These variations of *PTEN* expression are within physiological relevant levels where a modest decrease of 20% have been shown to increase cancer susceptibility^17^.

Our data is also consistent with previous RNA-sequencing studies showing induction of *C/EBP*β in BRAFi resistant cells^18^. However, these previous studies did not report *PTEN* or PTENP1-AS as candidate genes involved in BRAFi resistance, possibly due to the applied thresholds where genes were required to have a fold change ≥ 2. Based on our studies, both *PTEN* and PTEN1-AS are likely to be excluded when these thresholds are applied.

Several PTENP1-encoded lncRNAs with opposing functions have been described. While the miRNA sponge model suggests concordant expression of *PTEN* and PTENP1-S, we have not observed this phenomenon in the present study^10^. Instead, the PTENP1-AS transcript appears to be the dominant regulator of *PTEN* expression under these conditions. However, this does not exclude that miRNA sponging could take place as well. A regulatory mechanism, where translation as well as transcription of *PTEN* is regulated through lncRNAs encoded by the PTENP1 locus is still a plausible scenario. Subtle variations of *PTEN* expression have previously been reported to be associated with cancer susceptibility^17^, suggesting that strict and highly ordered regulation of *PTEN* expression is crucial for evading carcinogenesis. It is therefore possible that *PTEN* is regulated at several different levels. For example, transcriptional regulation by the PTENP1-AS may be the dominant regulator of *PTEN*, while fine adjustment may be taking place by miRNA sponging and post-transcriptional effects. Deletions of PTENP1 have been reported in melanoma cell lines and tissues, supporting that suppression of *PTEN* can occur also at the post-transcriptional level^19^. Additionally, PTENP1-S may also be involved in sponging of miRNAs related to other mRNAs beyond *PTEN*^20^.

Notably, this study also shows that the suppression of *PTEN* in BRAFi resistant cells may be reversible through targeting of *EZH2, DNMT3A* as well as PTENP1-AS. The, individual knockdown of *EZH2* or *DNMT3A* alone did not re-activate the expression of *PTEN* to the same extent as co-knockdown of these proteins (**Figures 3B-D**). We believe this is due to incomplete knockdown of *DNMT3A* and *EZH2* where some active protein complexes may still be present under these circumstances (**Figure S2**). An interaction between EZH2 and DNMT3a have been demonstrated^21^ and the combination of *EZH2* and *DNMT3A* targeting may in this case be more efficient in reducing the protein complexes more completely. Importantly, the induction of *PTEN* is more prominent in the A375PR1 cells than in the parental cells, indicative of a therapeutic window for re-activation of *PTEN* in BRAFi resistant melanoma cells. In support to this, we show that manipulation of the PTENP1-AS pathway re-sensitized the A375PR1 resistant cells to treatment with vemurafenib (**Figure 6A**).

We finally evaluated the expression of PTENP1-AS in regional lymph node metastases of stage III melanoma and found that high expression correlated with poor survival (**Figure 6C**). This indicates that PTENP1-AS is involved in melanoma tumor progression and is not only important during the development of resistance to BRAFi. Also, development of a particular type of resistance to vemurafenib may depend on the pre-existing factors/pathways in the tumors. High expression of PTENP1-AS could for example indicate that the tumor is less likely to respond to treatment. Moreover, loss of *PTEN* expression has previously been linked to metastasis^12^. Therefore, one may envision PTENP1-AS being involved in such inactivation and consequently enhance initiation of distant metastasis. Additional studies will be required in order to better understand the functional role and importance of PTENP1-AS in melanoma progression and drug resistance by taking patient samples pre- and post-treatment with BRAFi.

In summary, we have identified PTENP1-AS to be involved both in melanoma tumor progression and in development of resistance to vemurafenib. The data presented here show that PTENP1-AS is not only a promising target for re-activation of *PTEN*, but also a possible prognostic marker for clinical outcome in stage III melanoma.

## MATERIALS AND METHODS

### Cell cultures

A375 cells were purchased from ATCC (CRL-1619). A375, A375PR1, A375VR3 and A375VR4 cell lines were cultured in 5% CO2 at 37 °C in MEM supplemented with 10% heat-inactivated FBS, 2 mM glutamine, 0.1mM non-essential amino acids, 1mM sodium pyruvate, 50 μg/ml of streptomycin and 50 μg/ml of penicillin.

### RNA extraction and cDNA

RNA was extracted using the RNA NucleoSpin II kit (Macherey-Nagel) and treated with DNase (Ambion Turbo DNA-free, Life Technologies). DNase treated RNA (∼500 ng) was used for the generation of cDNAs using MuMLV (Life Technologies) and a mixture of oligo(dT)15 with nanomers.

### Semi-qRTPCR

PCR was performed by using the KAPA2G FAST mix (Kapa Biosystems) according to the manufacturer’s recommendations and by using the corresponding oligos in supplementary Table 1.

### qRTPCR

qRTPCR was performed by using the KAPA 2G SYBR Green (Kapa Biosystems) on the Applied Biosystems 7900HT or the BioRad CFX96 Touch platform with the following cycling conditions: 95 °C for 3 min, 95 °C for 3 s, 60 °C for 30 s. The corresponding oligos for each target is specified in supplementary Table 1.

### siRNAs and gapmers

siRNAs and gapmer antisense oligos (ASO) were ordered from the respective manufacturers (supplementary Table 1) and transfected by using lipofectamine 2000 (Life Technologies) according to the manufacturer’s recommendations. A final concentration of 10-40 nM was used for the siRNAs and gapmer ASOs and PLUS Reagent (Life Technologies) added for transfections of gapmers.

### Protein analysis

Samples were lysed in 50 mM Tris-HCl, pH 7.4, 1% NP-40, 150 mM NaCl, 1 mM EDTA, 1% glycerol, 100 μM vanadate, protease inhibitor cocktail and PhosSTOP (Roche Diagnostics GmbH). For the analysis of histones, the samples were subjected to sonication using a Bioruptor Sonicator (Diagenode), 30sec ON, 30sec OFF (setting = high) for a total of 6 cycles. Lysates were subjected to SDS-PAGE using 4-12% acrylamide gels (Life Technologies) and transferred to PVDF membranes using the iBlot system (Life Technologies). The proteins were detected by western blot analysis by using an enhanced chemiluminescence system (Western Lightning– ECL, PerkinElmer). Antibodies used were specific for PTEN (Cell Signaling, cat. no. 9552, 1:1,000), AKT (Cell Signaling, cat. no. 9272, 1:1,000), phospho-AKT (Cell signaling, cat. no. 4060S, 1:1,000) and β-actin (Sigma-Aldrich, cat. no. A5441, 1:5,000).

### ChIP of EZH2, H3K27me3 and C/EBPβ

ChIP assays were performed as previously described^8^. Briefly, the ChIP assay Kit (Upstate/Millipore) was used by crosslinking the cells in 1% formaldehyde for 10 minutes, quenched in 0.125M Glycine for 5 minutes and lysed according to the manufacturer’s recommendations. The samples/nuclei were sonicated with a Bioruptor Sonicator (Diagenode) at 30sec ON, 30sec OFF (setting = high) for a total of 18 cycles. The water was replaced with ice-cold water after every sixth cycle. The samples were diluted 1:10 in ChIP dilution buffer, pre-cleared and incubated over night with the appropriate antibody. Salmon sperm DNA/Protein A–agarose (Upstate/Millipore) was used to pull down the antibody. The DNA was eluted in elution buffer (1% SDS, 100 mM NaHCO3), followed by reverse cross-linking at 65°C over night. The samples were RNase-A and protease-K treated and finally eluted using the Qiagen PCR purification kit (Qiagen). The following antibodies were used (4 μg/sample): H3K27me3 (Upstate/Millipore, cat. no. 17-622), EZH2 (Upstate/Millipore, cat. no. 07-689) and C/EBPβ (Santa Cruz Biotechnology cat. No sc-150).

### Assessment of methylated DNA

The enzyme McrBc (New England Biolabs) was used to cleave methylated DNA. Briefly, 200 ng DNA was digested with McrBc at 37°C overnight and heat inactivated the next day at 65° for 1 h. Samples were run on a qPCR and standardized to uncut input. Delta CT values were converted to fold-change values and the ratio between A375 PR1/A375 calculated.

### PI-annexin V staining

The cells were harvested, washed twice in PBS and resuspended in 100 μL annexin V incubation buffer (10 mM HEPES/NaOH, pH 7.4, 140 mM NaCl, 5 mM CaCl2) containing 1% annexin V FLOUS (Roche Molecular Biochemicals) and 500 μg/μl PI stain. The samples were incubated for 15min at room temperature followed by adding 400 μL of ice-cold annexin V incubation buffer and subsequently analyzed on a cytometry machine.

### Patient samples

Samples from stage III melanoma patients with first regional lymph node metastases that had not received any systemic treatment were collected. The RNA was extracted using the Qiagen RNeasy mini kit (Qiagen). The RNA was DNase treated on column according to the manufacturer’s recommendations. The expression of PTENP1-AS was analyzed using qRTPCR and standardized to *beta-actin* (dCt) by using the corresponding oligos in Supplementary Table 1. ddCt was calculated by using the mean Ct value of all samples. High expression of PTENP1-AS (Figure 7B) was defined as Ct < 30.

### Statistics

Two tailed Student’s T-test was used to determine statistical significance. Error bars represent the standard error of the mean. Statistical significance of the melanoma patient samples in figure 7B is evaluated using log-rank test. Throughout the paper, * P < 0.05; ** P < 0.01; *** P < 0.005.

## ACKNOWLEDGEMENTS

Professor Dan Grandér, who was the grant holder and the senior author of this project at the Department of Oncology & Pathology, passed away in October 2017. The authors are deeply sorrowed by his tragic death and dedicate this study to his memory.

## DECLATATION OF INTEREST STATEMENT

The authors declare no conflicts of interest.

## AUTHOR CONTRIBUTION

L.V, A.A, I.S, and P.J performed experiments. A.S.R and O.S provided general laboratory support for the research project. A.P and S.K designed gapmer antisense oligonucleotide. J.H, S.E.B, C.I, G.J, H.O, M.F.S and R.T provided and prepared clinical samples. L.V, D.G and P.J wrote the manuscript with the support from A.A, S.E.B, J.H and K.P.T. D.G and P.J supervised the project.

## COMPETING INTEREST

The authors declare no competing interests.

### Ethical approval and consent to participate

This study has been approved by the Research Ethics Committee of Karolinska Institutet (Dnr: 2006/1373-31/3) and the Regional Ethics Committee in Lund (Dnr 2013/101). All involved individuals have provided informed consent to be included in the study.

