## supplementary_data for "PTENP1-AS contributes to BRAF inhibitor resistance and is associated with adverse clinical outcome in stage III melanoma"

**SUPPLEMENTARY FIGURE LEGENDS**

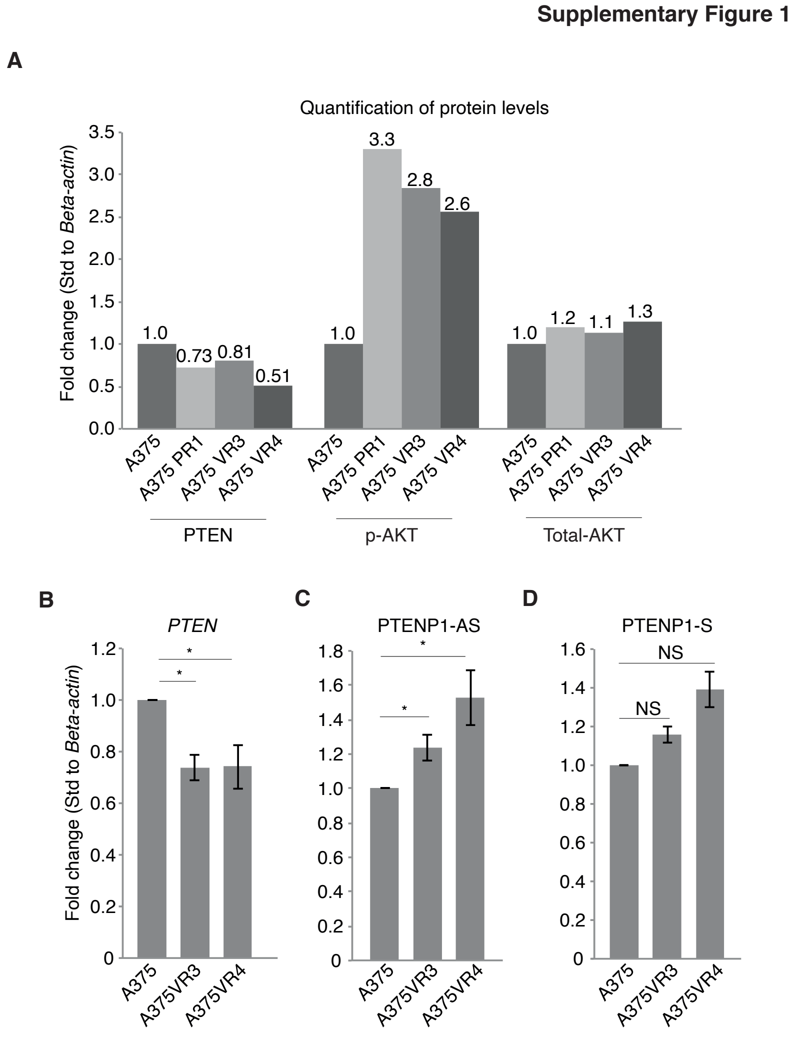

**Supplementary Figure 1** *RNA expression levels in BRAFi resistant A375 sublines*

**(A)** Quantification of western blot (Figure 1A) using the ImageJ software. **(B-D)** QRTPCR measurements of **(B)** *PTEN,* **(C)** PTENP1-AS and **(D)** PTENP1-S in BRAFi resistant A375 sublines relative A375 cells.

**Supplementary Figure 2**

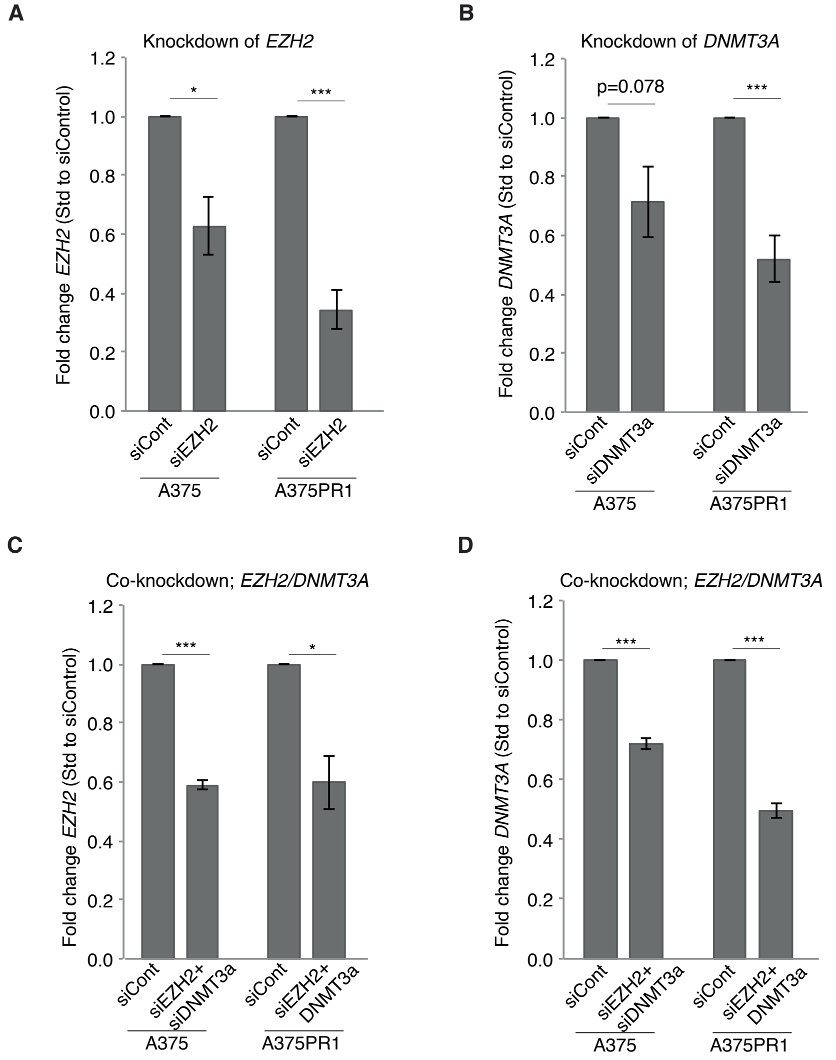

**Supplementary Figure 2** *Knockdown of EZH2 and DNMT3A*

**(A-B)** qRTPCR measurements of *EZH2* and *DNMT3a* in A375 and A375PR1 cells. (**C-D**) qRTPCR measurements of *EZH2* and *DNMT3A* in A375 and A375PR1 upon simultaneous knockdown *of EZH2/DNMT3A.*

**Supplementary Figure 3**

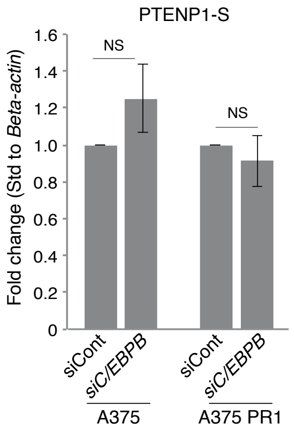

**Supplementary Figure 3**

**(A)** qRTPCR presenting the expression of PTENP1-S upon siRNA induced knockdown of *C/EBPβ* in A375 and A375 PR1 cells.

**SUPPLEMENTARY TABLE 1**

| | |
|---|---|
| **qRTPCR** | |
| B-actin F | AGGTCATCACCATTGGCAATGAG |
| B-actin R | CTTTGCGGATGTCCACGTCA |
| DNMT3a F | TTTGAGTTCTACCGCCTCCTGCAT |
| DNMT3a R | GTGCAGCTGACACTTCTTTGGCAT |
| EZH2 F | CAGTTTGTTGGCGGAAGCGTGTAA |
| EZH2 R | AGGATGTGCACAGGCTGTATCCTT |
| PTEN set I F (3'UTR) | AGA AAG CTT ACA GTT GGG CCC TGT |
| PTEN set I R (3'UTR) | GCC ACA GCA AAG AAT GGT GAT GCT |
| PTEN set II F (orf) | GGG ACG AAC TGG TGT AAT GAT ATG |
| PTEN set II R (orf) | CCA GAT GAT TCT TTA ACA GGT AGC TAT AA |
| PTENP1-AS F | CCTCACAGCGGCTCAACATTCAAA |
| PTENP1-AS R | AGGCTTCCAGGTTGGAAAGGAA |
| PTENP1-S F | AGTCACCTGTTAAGAAAATGAGAAGACAAA |
| PTENP1-S R | CTGTCCCTTATCAGATACATGACTTTCAA |
| C/EBPβ set 2 F | CGCGACAAGGCCAAGAT |
| C/EBPβ set 2 R | GCTGCTCCACCTTCTTCTG |
| **Detection of PTENP1-AS** | |
| PTENP1-AS F0 | AAG CCC ACG GCT TCC ACC TT |
| PTENP1-AS F2 | AGACGAAGAAGAAGCGAGAAACGC |
| PTENP1-AS F3 | CCTCACAGCGGCTCAACATTCAAA |

| | |
|---|---|
| PTENP1-AS R0 | GCT GCA ATA ATC AAC AGA GTG TGG |
| PTENP1-AS R3 | AGGCTTCCAGGTTGGAAAGGAA |

**ChIP**

| | |
|---|---|
| PTEN pro F | TGATGTGGCGGGACTCTTTATGC |
| PTEN pro R | TCACAGCGGCTCAACTCTCAAACT |

**siRNAs**

| | |
|---|---|
| EZH2 #1 | Cat#; SI02665166 (Qiagen) |
| EZH2 #2 | Cat#; SI00063959 (Qiagen) |
| DNMT3a #1 | Cat#; SI02665271 (Qiagen) |
| DNMT3a_11 | Cat#; SI02665278 (Qiagen) |

**PTENP1-AS siRNA**

| | |
|---|---|
| Sense | GACGAAGAAGAAGCGAGAAAC TT |
| Antisense | pho-GUUUCUCGCUUCUUCUUCGUC TT |

**DsiRNA**

| | |
|---|---|
| CEBPβ | AGUUGAUGCAAUCGGUUUAAACATG |
| | CAUGUUUAAACCGAUUGCAUCAACUUC |

**GAPMERS**

| | |
|---|---|
| Gapmer Control | CGAATAGTTAGTAGCG |
| Gapmer PTENP1-AS | CGTACAGATAAGAGGATTA |
